# Dynamics of Reactive Oxygen Species and Nitrogen Cycling in Soils as a Mechanism for Volatile Reactive Nitrogen Oxide Production

**DOI:** 10.1101/2024.11.29.625130

**Authors:** Megan L. Purchase, Jonathan D. Raff, Alexander G. Gale, Alannah Vaughan, Gary D. Bending, Ryan M. Mushinski

**Affiliations:** School of Life Sciences, University of Warwick, Coventry CV4 7AL, United Kingdom; Paul O’Neill School of Public and Environmental Affairs and Department of Chemistry, Indiana University, Bloomington, USA

## Abstract

Heterotrophic bacteria and fungi are responsible for the decomposition of organic matter in soil. During this process, reactive oxygen species (ROS) can be produced directly and indirectly as extracellular byproducts of respiration. It is well known that nitrogen (N) cycle processes lead to the formation of volatile reactive nitrogen oxides (NO_y_), a group of climate-active gases that contribute to atmospheric chemistry and negatively impact human health. Primary microbial sources of NO_y_ include ammonia- oxidising bacteria and denitrifying bacteria and fungi. Despite soil being a significant source of global NO_y_ emissions, studies to date have primarily focused on N emissions from agricultural soils and there remains a large scope for investigating mechanisms of soil NO_y_ production in a wider context in order to better constrain terrestrial and climate process models. Here, we propose a potential microbial mechanism involving ROS that could influence the production of NO_y_ from soil. We utilised metagenomics and metatranscriptomics alongside continuous gas flux measurements and analysis of soil properties to evaluate the connection between microbial ROS and soil-sourced NO_y_. Our findings suggest that more NO_y_, particularly nitric oxide (NO), is produced in the presence of increased abundance of ammonia-oxidising and denitrifying taxa. NO can be lost to the environment or reacts with superoxide, an ROS produced via the enzymatic activity of soil organic matter (SOM) decomposers. This reaction produces peroxynitrite (ONOO^-^), which we have demonstrated to enhance nitrogen dioxide (NO_2_) emissions from soil. We have shown that the extent of NO_y_ production through this pathway may be dependent on SOM composition, and the associated variability in carbon and nitrogen content.

## 1. Introduction

Complex chemical and biological dynamics at the biosphere-atmosphere interface lead to the formation and emission of volatile reactive nitrogen oxides (NO_y_), a group that includes NO_x_ (nitric oxide [NO] + nitrogen dioxide [NO_2_]) and NO_z_ (nitrous acid [HONO] + nitric acid [HNO_3_] + nitrogen trioxide [NO_3_] + …). NO_y_ species contribute to atmospheric pollution, climate change, and reducing air quality. Soil-sourced NO_y_ is a significant contributor to total NO_y_ emissions^1,2^, yet there is a lack of mechanistic data relating to the origin of these gases in soil systems, resulting in uncertainty in biogeochemical and atmospheric models. The nitrogen (N) cycle intersects with various other concurrent biogeochemical processes, leading to a dynamic pool of compounds that may react under thermodynamically favourable conditions.

### 1.1 Mechanisms of Soil N Cycling

N can enter soil naturally through biological N-fixation and artificially through fertilisation and atmospheric N-deposition.^3^ This creates a pool of reactive N (N_r_), typically in the forms of ammonia/ammonium (NH_3_/NH_4+_) or nitrate (NO_3-_). A large proportion of this N_r_ is immobilised by plants and microbes for synthesis of N-dependent macromolecules, such as proteins. The residual N_r_ pool can undergo microbial oxidative and reductive catalysis^4^ as a means of energy generation for chemoautotrophic microbes with NO_y_ as products and by-products. Rates of organic matter decomposition and N-cycling are highly variable within soils, with the availability of soil organic matter (SOM) being a primary mediator.^5^ SOM can be subcategorised into particulate organic matter (POM) and mineral associated organic matter (MAOM) based on particle size or density, with different routes of formation, persistence times, and functions.^6^ SOM content and composition is important for soil fertility and structure, and is dependent on organic matter input and decomposition rate, and SOM mineralisation rate. ^7–9^ Dominance of either the MAOM or POM fraction in soil defines the N availability and ratio of carbon (C) to N (C:N), which will then determine rates of microbial activity.^10^ The parent material and texture of soils can aid or hinder the accumulation of SOM through physical and chemical protection from microbial acquisition; sandy soils generally contain less SOM than clayey soils.^11,12^ It has been reported that soils with low C:N show an increase in N-cycle microbial biomass, as well as increased rates of mineralisation and nitrification.^13^ Despite the biogeochemical significance of SOM composition, studies directly linking SOM to soil- sourced NO_y_ are lacking.^14,15^

### 1.2 Drivers of NOy Emissions

Land-use change from natural ecosystems, such as woodland and grassland, to human-influenced agricultural and urban areas is an increasingly prominent agent of change to soil properties including SOM composition, microbial communities, and nutrient cycling.^16,17^ Our previous work has established soil from different land-use types exhibit significant differences in N-cycle process rates, and NO_y_ fluxes.^18^ Soils of different land-use types are subject to various chemical and physical disturbances as a result of human activity such as fertiliser application and fossil fuel combustion, and natural processes such as mineral weathering and the addition of organic matter to soils via leaf litter of variable composition and quality. Globally, increased N deposition to both aquatic and terrestrial systems is closely associated with increased anthropogenic activity.^19^ Excessive combustion of fossil fuels, and intensification of modern agricultural practices using inorganic N fertilisers, means that large quantities of N are lost to the environment – creating a situation where the maximum amount of N immobilisation is exceeded and processes that lead to NO_y_ are exacerbated. Research to date has established a positive relationship between N deposition and increased emissions of nitrous oxide (N_2_O) from soils^20–22^, however studies on the effects of N deposition on soil-sourced NO_y_ are far fewer. N deposition to soil leads to changes in microbial biomass N and soil C:N, soil microbial growth and diversity, and the dynamics of N-cycling.^19^ Human activity that leads to N deposition, such as combustion of fuels, often also leads to deposition of heavy metals to soils, altering soil pH and leading to acidification and subsequently affecting microbial community structure and function.^23,24^ NO_y_ is also produced abiotically from the intermediates hydroxylamine (NH_2_OH) and NO_2-_ from the microbial N-cycle. These reactions depend on concentrations of minerals, particularly those containing iron, as well as soil pH and oxygen (O_2_) availability.^25,26^

### 1.3 Identifying Concurrent Chemistry of Soil ROS and N cycling

Once released into soil, N-cycling intermediates are exposed to a number of oxidants including, reactive oxygen species (ROS) such as superoxide (O_2•−_), hydroxyl (•OH) radical, and hydrogen peroxide (H_2_O_2_). Microbial respiration is a primary source of ROS in soils as microbes decompose SOM. NADPH oxidases (NOX) are a significant source of ROS which can be involved in the breakdown of SOM.^27–29^ In animal and plant physiology, ROS and N species interact to control many biological processes – mostly controlled by the activity of superoxide dismutase (SOD).^30^ Of particular interest is O_2•−_ which reacts readily with NO to form peroxynitrite (ONOO^-^).^31^ ONOO^-^ can subsequently interact chemically with soil-prevalent compounds, including CO_2_, to form NO_2_, NO_3_ˉ, and HONO.^32^ It is reasonable to hypothesise that such processes abound in the soil environment, although to our knowledge this has not been previously investigated. The objective of this study is to explore unrecognised connections between the ROS and N cycles that impact soil-derived emissions of NO_y_ to the atmosphere. We did this by examining soil from the land-use type and urbanisation gradient of the U.K., which provided a robust dataset with a wide range of soil properties. Coupling metagenomic and metatranscriptomic approaches provided mechanistic data on soil ROS and N cycling that help to constrain the most important drivers of biogenic soil-sourced NO_y_ species that are often overlooked in the chemical transport models used to study climate change and air pollution.

## 2. Methods

### 2.1 Soil Sampling

Soils were collected in October 2022 from three land-use types: agricultural, woodland, and built. Built soils are defined as those <1 m from a human-made structure. Each land-use was sampled at University of Warwick Stratford Innovation Campus, and Coventry City, corresponding to low (<1000) and high (>400,000) human populations (**Fig. S1**). Three replicates were taken of each land-use at each location, giving a total of 18 samples. One intact core (0-10 cm depth) per land-use type was taken for each location.

### 2.2 Analysis of Soil Microbiomes

RNA and DNA were extracted from 2 g fresh soil according to the Qiagen RNeasy PowerSoil Total RNA Kit and RNeasy PowerSoil DNA Elution Kit. Sequencing was conducted by the NERC Environmental Omics Facility (NEOF) at the Centre for Genomic Research, University of Liverpool. Submitted total RNA was used to prepare a dual-indexed, strand-specific RNASeq library using the QIAseq FastSelect rRNA depletion and Ultra II Directional RNA library preparation kits. Submitted gDNA was used to prepare an Illumina fragment library using the NEBNext Ultra II FS Kit. RNA and DNA were sequenced using Illumina NovaSeq paired-end, 2 x 150bp. Analysis of DNA and RNA sequencing data was carried out on a system equipped with Dell PowerEdge R640 compute nodes, each with 2 x Intel Xeon Platinum 8268 2.9 GHz 24-core processors, 48 cores per node, 1536 DDR4-2933 RAM per node, and 32 GB RAM per core. Paired-end reads from DNA and RNA sequencing were first merged using Paired-End reAd mergeR (PEAR) (v0.9.8).^33^ PEAR implements normalisation methods of quality score adjustment, consensus base calling, and pre-merging quality trimming to generate high-confidence merged reads. Reads were aligned against the NCBI-nr protein database using Double Index AlignMent Of Next-generation sequencing Data (DIAMOND), which included adapter trimming and quality filtering, and length and coverage normalisation.^34^ Rarefaction curves were generated to confirm adequate sequencing depth. Reads and alignments were mapped to taxonomic and functional classes using the meganizer tool from MEtaGenomic ANalysis 6 Ultimate Edition (MEGAN6).^35^ MEGAN6 was used for further analysis.

### 2.3 Quantification of Soil NO_y_ Gas Fluxes

Fluxes of potential total NO_y_, NO, and NO_2_ + NO_z_ were analysed from intact soil core microcosms as in Purchase *et al.*^18^ with a chemiluminescence technique using a Teledyne T200U-NO_y_ analyser with an external NO_y_ converter (Teledyne API, CA, USA). The lower detectable limit for the measurement of NO with this instrument was measured to be 100 ppt for 70 s averages. Flux calculations are detailed in **Method S1** of the supporting information.

### 2.4 Quantification of Peroxynitrite Decomposition

To assess whether the thermodynamic product of NO and O_2•−_ (e.g., ONOO^-^) could lead to NO_y_ production, soil microcosms were amended with varying levels of ONOO^-^. Synthesis of ONOO^-^ followed the protocol devised by Robinson and Beckman^36^. Briefly, hydrogen peroxide acidified with hydrochloric acid was reacted with sodium nitrite to produce peroxynitrous acid. The reaction was quenched with an excess of sodium hydroxide to produce the ONOO^-^ anion in solution at approximately 180 mM concentration. In one set of experiments, 15 mL of dilutions of ONOO^-^ solution (0.32, 0.16, 0.08 µM) in deionised (DI) H_2_O, and a blank of only DI H_2_O were added to 200 g soil in four microcosms, and fluxes of NO_2_ were measured over 1 h as in **Method 2.3**. Microcosm design is detailed in Purchase *et al*..^18^ In a second set of experiments, we compared NO, NO_2_, and NO_z_ head space accumulation in soil slurries with and without added ONOŌ (0.01 µM). Specifically, 5 g soil (agricultural, grassland, and woodland) were added to 125-mL Wheaton bottles, suspended in 15 mL water or 0.01 µM ONOŌ, and capped with butyl septa. Bottles were shaken at 150 rpm at 25 °C in the dark for 8 h. Headspace samples for NO, NO_2_, and NO_z_ were collected through butyl stoppers and analysed immediately with a chemiluminescence detector.

### 2.5 Soil Organic Matter Composition

Soils were first passed through a 2 mm sieve, then dried at 65 °C for 48 h. To disperse the soils, oven dried samples were shaken in dilute 0.5% sodium hexametaphosphate and 3 mm glass beads for 18 h. To separate the samples into MAOM and POM fractions, soils were rinsed onto a 53 µm sieve. The fraction that passed through was collected as MAOM and the remaining fraction was collected as POM, as described in Cotrufo *et al..*^10^ Although POM and MAOM fractions cannot be definitively distinguished by size fractionation, previous studies have found this method to be as effective as others, such as density fractionation.^37,38^

### 2.6 Soil Texture Analysis

Soils were dried at 80 °C for 24 h. To disperse the soils, oven dried samples were shaken in dilute 0.5% sodium hexametaphosphate and DI H_2_O. Sample mixtures were transferred to 1 L graduated sedimentation cylinders and DI H_2_O was used to bring the volume to 1 L. The contents were thoroughly vertically mixed using a plunger. Hydrometer and temperature readings were taken immediately after mixing, and again after the cylinder had been left undisturbed for 2 h. Hydrometer readings were corrected for temperature by subtracting 0.36 g L^-1^ from the hydrometer reading for each degree below 20 °C, and the corrected blank reading was subtracted from corrected sample readings. Silt, clay, and sand fractions (% L^-1^) were calculated from corrected hydrometer readings (see **Method S2** of the supporting information).

### 2.7 Quantification of Soil Total Carbon and Nitrogen Content

Soils were dried at 60 °C for 24 h. For each sample, 1 g soil was weighed into a 2 mL stainless steel grinding tube with two stainless steel 5 mm ball bearings. Tubes were sealed with silicon bungs. A FastPrep-24™ Classic bead beating grinder (MP Biomedicals, CA, USA) was used to grind the samples in 1 min increments 12 times. Samples were analysed for total C and N content using a vario PYRO cube^®^ elemental analyzer (Elementar UK Ltd.). Total soil samples and individual MAOM and POM fractions were analysed.

### 2.8 Quantification of Soil pH and Soil Processes

10 g subsamples of soil were air-dried for 48 h then suspended in 20 mL of 0.01 M calcium chloride (CaCl_2_) for pH measurement using a pH meter. Microbial soil respiration (mg-CO_2_ kg-soil^-1^ d^-1^) was determined using the WTW Xylem Analytics OxiTop^®^-IDS measurement system according to ISO 16072 (**Method S3**). 100 g soil was incubated for 7 d and the change in pressure each day was recorded. 3 g soda lime was used as an adsorbent for CO_2_ produced by microorganisms during the sampling period. A segmented flow analyser (Seal Analytical Ltd, Wrexham, UK) was used to analyse concentrations of NO_3-_ + NO_2-_(AutoAnalyser Method G-109-94, ISO 13395) and NH_4+_ (AutoAnalyser Method G 102-93, ISO 11732), as in Purchase *et al..*^18^ Concentrations were converted from a mass nutrient per unit volume to a mass nutrient per mass dry soil. Net nitrification, net ammonification and net total N mineralisation rates were calculated as in **Method S4**.

### 2.9 Statistical Analyses

Measurements of potential mean NO_y_ fluxes were made from 6 samples, with 5 datapoints per sample. All other analyses were carried out on 18 samples. Statistical tests were performed using R (v4.1.2).^39^ Shapiro-Wilk tests were carried out to test for normality of all datasets. To assess the relationship between measured variables and land-use types or locations of sample sites, Kruskal-Wallis rank sum tests (KW) were used. To assess relationships between measured variables, Spearman’s rank correlation coefficient tests (SRCC) were used. Significant differences were inferred when *p* < 0.05. *P*-values from Kruskal-Wallis tests were corrected for multiple comparisons with a Dunn’s test using the false discovery rate with the Benjamini-Hochberg method using R package *FSA_0.9.4*. Non-metric multidimensional scaling analysis (NMDS) and analysis of similarity (ANOSIM) were carried out using R packages *vegan_2.6-4* and *plyr_1.8.8*. Partial Least Squares Path Models (PLS-PM) and mixed effects models were created using R packages *lme4_1.1-35.3, plspm_0.5.1* and *sjPlot_2.8.16*. Model validation for PLS-PM was performed using bootstrapping (1000 iterations) to obtain confidence intervals for path coefficients. Goodness-of-fit was assessed using standardised root mean square residual and normal fit index. A priori power analysis using G*Power_3.1^40^ indicated that our sample size (n=18) provided 95% power to detect large effects (f^2^ > 0.35) at α = 0.05.

## 3 Results

### 3.1 Metagenomics and Metatranscriptomics

For metagenomic (DNA) sequencing, a total of 347,641,824 reads were obtained from 18 samples. The average reads assigned per sample was 5,607,470, ∼29%. Alignment with the NCBI-nr database identified 96 bacterial orders, 1 archaeal order, and 12 fungal orders. For metatranscriptomic (RNA) sequencing, a total of 332,969,312 reads were obtained from 18 samples. The average reads assigned per sample was 12,444,231, ∼20%. Alignment with the NCBI-nr database identified 97 bacterial orders, 2 archaeal orders, and 49 fungal orders. Subsequent analysis was carried out at the order level. Indicator species analysis (ISA) reveals that abundances of 67.2% of microbial orders were significantly different between DNA and RNA. Simpson’s reciprocal diversity index showed significantly greater alpha diversity from RNA compared to DNA. From non-metric multidimensional scaling (NMDS) and analysis of similarity (ANOSIM), there were no significant differences in microbial community composition between land-use types for DNA or locations for DNA or RNA. However, significant differences in the composition of the microbial community were indicated between land-use types of RNA samples (R = 0.2428, *p* < 0.05) (**Fig. 1A & 1B**).

**Figure 1.**
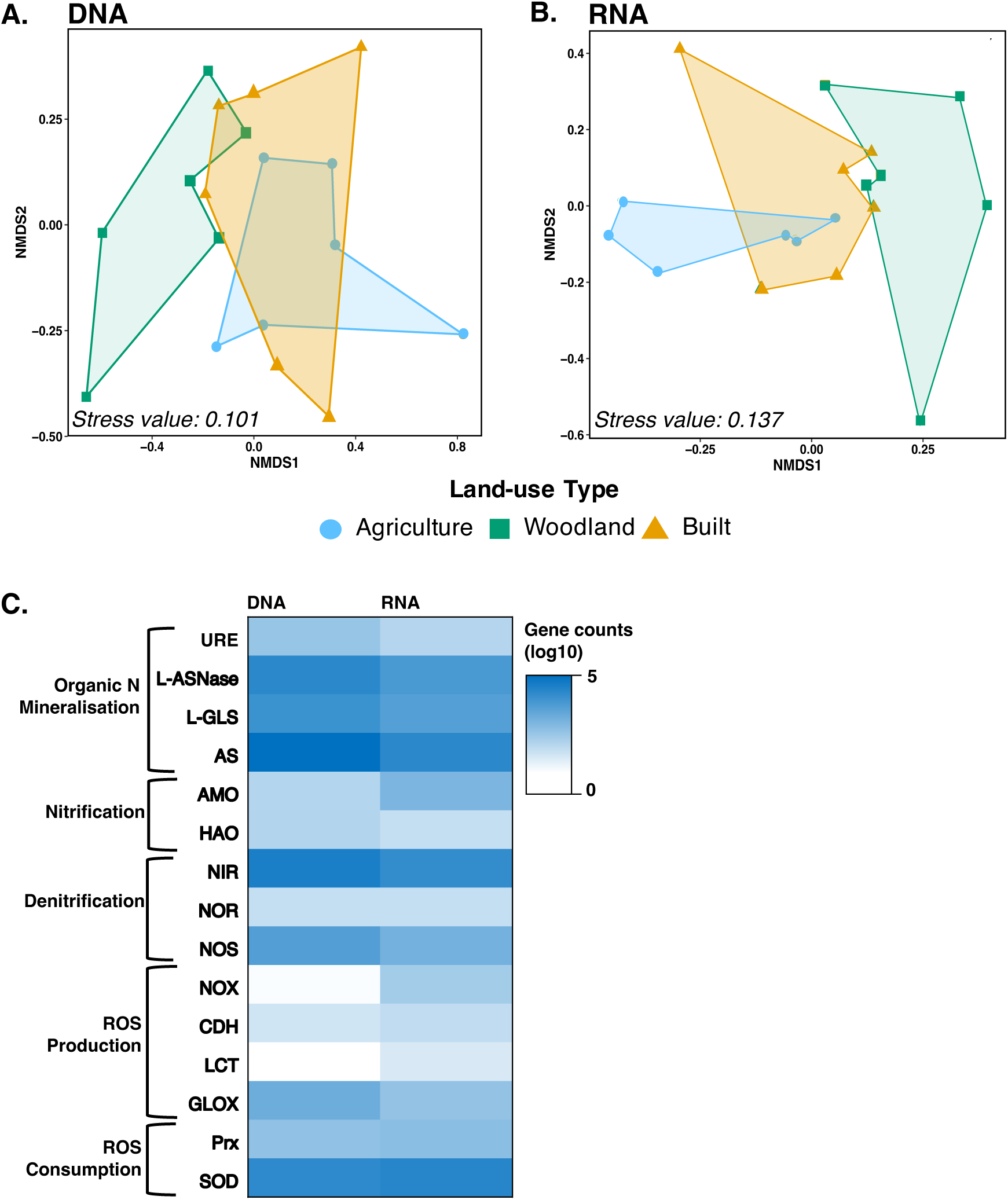
**(A)** Non-metric Multidimensional Scaling (NMDS) analysis of microbial communities from RNA sequencing between land-use types (“agriculture”, “woodland”, “built”). **(B)** NMDS analysis of microbial communities from DNA sequencing between land-use types. **(C)** Heat map of log10 gene counts from DNA and RNA sequencing of N cycling and ROS cycling microbes. *N* = 18.

Following functional assignment using the KEGG database within MEGAN6, abundances of specific N cycle and ROS cycle genes of interest were analysed (**Fig. 1C**). Abundance of microbes with genes that encode N-cycle genes L-glutaminase (L- GLS), urease (URE), ammonia monoxygenase (AMO), and nitrite reductase (NIR), and the ROS-cycle gene glyoxal oxidase (GLOX) were significantly associated with land-use type, and all were most abundant in woodland soils (KW, with Dunn’s test, *p* < 0.05, for all). Metatranscriptomes of L-GLS and NOX were significantly associated with land-use type, both being more abundant in built soils (KW, with Dunn’s test, *p* < 0.05, for both). Metatranscriptomes of NIR and SOD were significantly associated with sample location, with highest activity of both genes in soils from locations with high human populations. (KW with Dunn’s test, *p* < 0.05, for both).

### 3.2 NO_y_ Fluxes

Potential mean fluxes of NO (*F_NO_*) were significantly associated with land-use type, with highest fluxes from woodland samples (KW with Dunn’s test, *p* < 0.01). Potential mean fluxes of NO_2_ + NO_z_ (*F_NO2+NOz_*) were also significantly associated with land-use type (KW with Dunn’s test, *p* < 0.05). *F_NO2+NOz_* from agricultural and built soils were significantly higher than those from woodland soils (**Table S1**). Metatranscriptomes of N-cycle genes (URE, L-asparaginase (L-ASNase), L-GLS, amidase (AS), AMO, hydroxylamine oxidoreductase (HAO), NIR, nitric oxide reductase (NOR), and nitrous oxide reductase (NOS)), and ROS-cycling genes (NOX and cellobiose dehydrogenase (CDH)) were significantly positively correlated with *F_NOy_* (SRCC, *p* < 0.001, for all) (**Fig. 2**). The metatranscriptome of lactase (LCT) was significantly negatively correlated with *F_NOy_* (SRCC, *p* < 0.001). The metatranscriptomes of NOS and GLOX was significantly positively correlated with *F_NO2NOz_* (SRCC, *p* < 0.05, for both).

**Figure 2.**
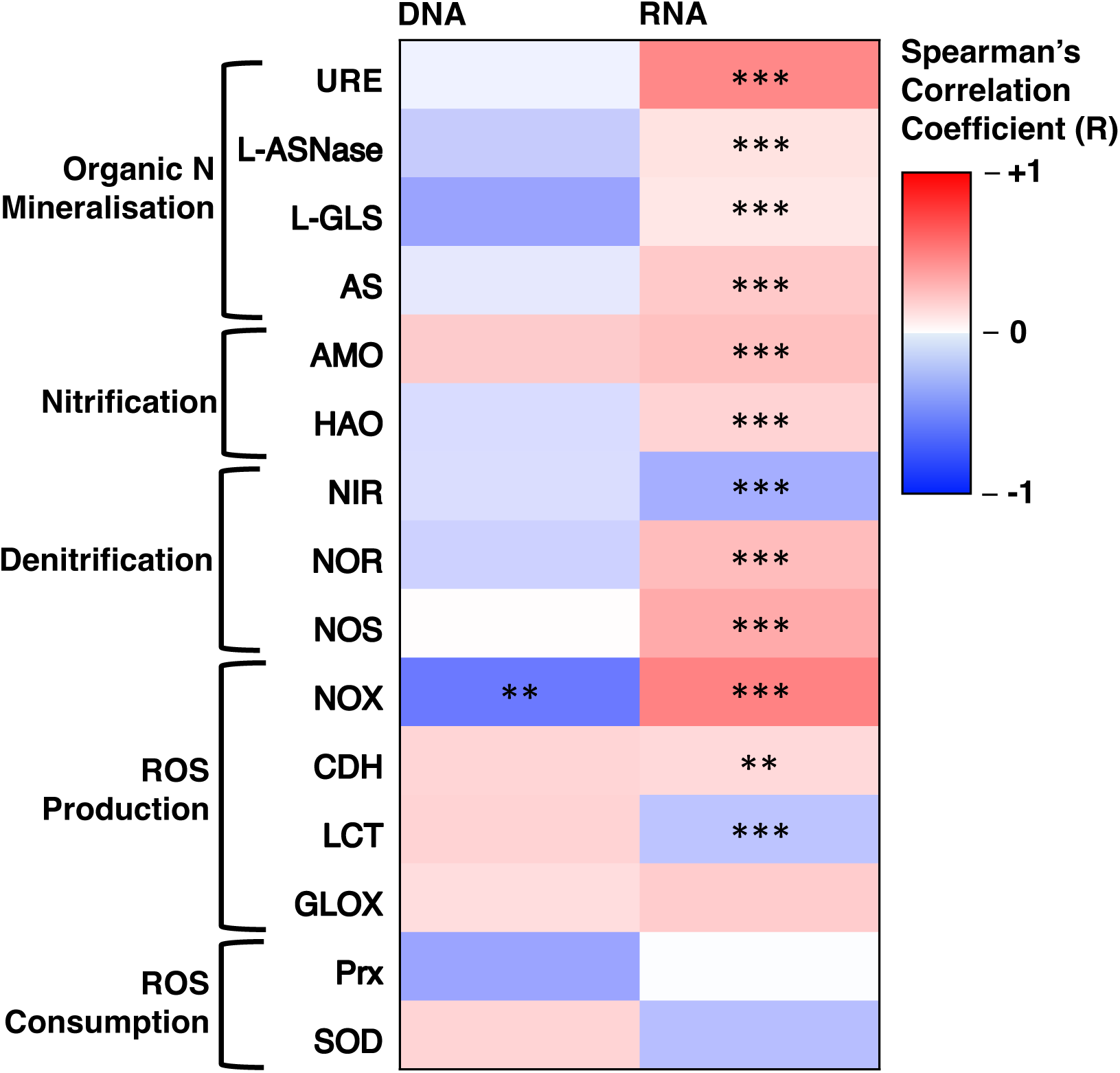
Heat map of Spearman’s Correlation Coefficients (R) testing the correlation of *F_NOy_* against metagenomic (DNA) and metatranscriptomic (RNA) data for genes associated with N cycling and ROS cycling. Microbial communities were quantified using Illumina NovaSeq. *N* = 18.

Abundance of microbes with genes that encode AMO were significantly positively correlated with *F_NO_* (SRCC, *p* < 0.05). Abundance of microbes with genes that encode NOX and L-GLS were significantly negatively correlated with *F_NOy_* and *F_NO2NOz_*, respectively (SRCC, *p* < 0.05, for both).

### 3.3 Quantification of Peroxynitrite Dynamics in Soil

For the soil chamber measurements, *F_NO2+NOz_* from soils in the first hour of measurements were significantly higher from soils with 0.08 µM and 0.32 µM ONOO^-^ addition compared to soils with DI H_2_O addition (KW, *p* < 0.01, for both) (**Fig. 3A**). For the soil slurry experiment, we observed a 60% increase in NO_2_ (one-way ANOVA, *p* < 0.01) and 47% increase in NO_z_ accumulation (one-way ANOVA, *p* < 0.05), respectively, in soils amended with ONOŌ relative to samples left unamended. (**Fig 3B**).

**Figure 3.**
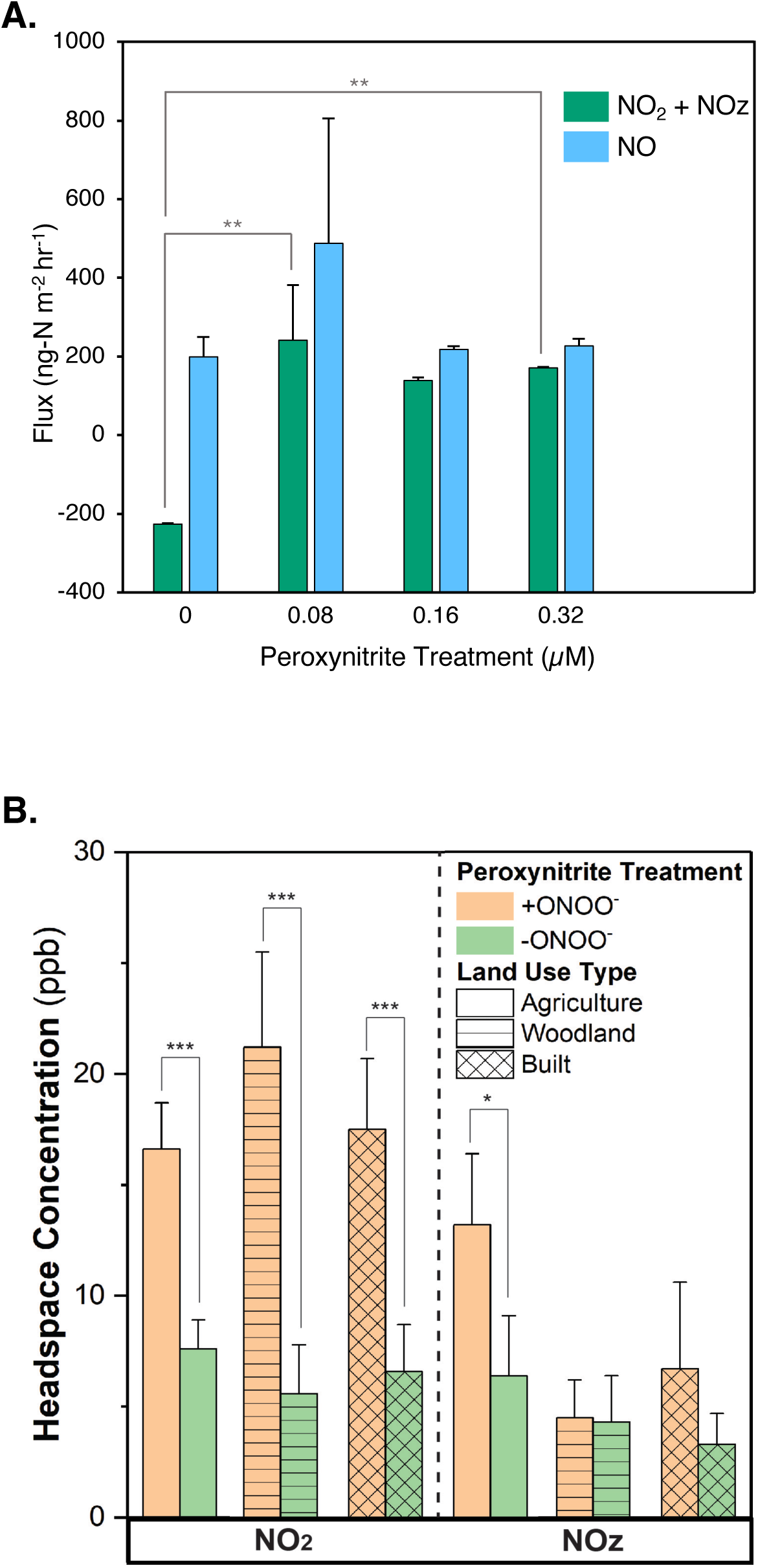
(A) Effect of peroxynitrite addition (µM) on potential 1 hr fluxes of NO and NO_2_+NO_z_ from soil. *N* = 4. **(B)** NO_y_ headspace accumulation in response to peroxynitrite addition (µM). Fluxes were measured with a chemiluminescence technique using a Teledyne T200U instrument. Significance is indicated by * = *p* < 0.05, ** = *p* < 0.01, *** = *p* < 0.001.

### 3.4 Soil Organic Matter Composition, Particle Size Distribution, and C and N content

SOM composition differed significantly between low human population and high human population locations (KW with Dunn’s test, *p* < 0.05), with higher MAOM:POM ratio in samples from high human population locations. There was a significant positive correlation between MAOM:POM and *F_NO_* (SRCC, *p* < 0.05) (**Fig. S2A**). There was also a significant positive correlation between MAOM:POM and soil moisture content (SRCC, *p* < 0.01). Mixed effects modelling indicates that the metatranscriptomes of N-cycle genes NOS, AS and L-GLS were positively associated with MAOM:POM, whereas L-ASNase was negatively associated (**Fig. 4**). The ROS-cycle genes we investigated were all negatively associated with MAOM:POM, particularly SOD. Soil texture classifications can be found in **Table S2** of the supporting information. There was no significant difference in clay, silt, or sand % L^-1^ between land-use types. Clay % was significantly positively associated with high human population locations (KW; *p* < 0.05) (**Fig. S2B**). Sand % was significantly positively associated with low human population locations (KW; *p* < 0.05). The C:N ratios of soil ranged from 8.43 to 19.12. There were no significant differences in total soil C:N between locations or land-use types. Total soil C:N was also not significantly associated with any potential gas flux measurements or soil moisture content. From mixed effects models, total soil C:N was positively associated with metatranscriptomes of N-cycle genes AS, and L-ASNase. Total soil C:N was also positively associated with metatranscriptomes of ROS-producing enzymes SOD, and LCT. C:N from MAOM fractions ranged from 9.85 to 41.48. C:N from POM fractions ranged from 8.44 to 14.32. C:N was significantly higher from MAOM fractions than POM fractions (KW; *p* < 0.001).

**Figure 4.**
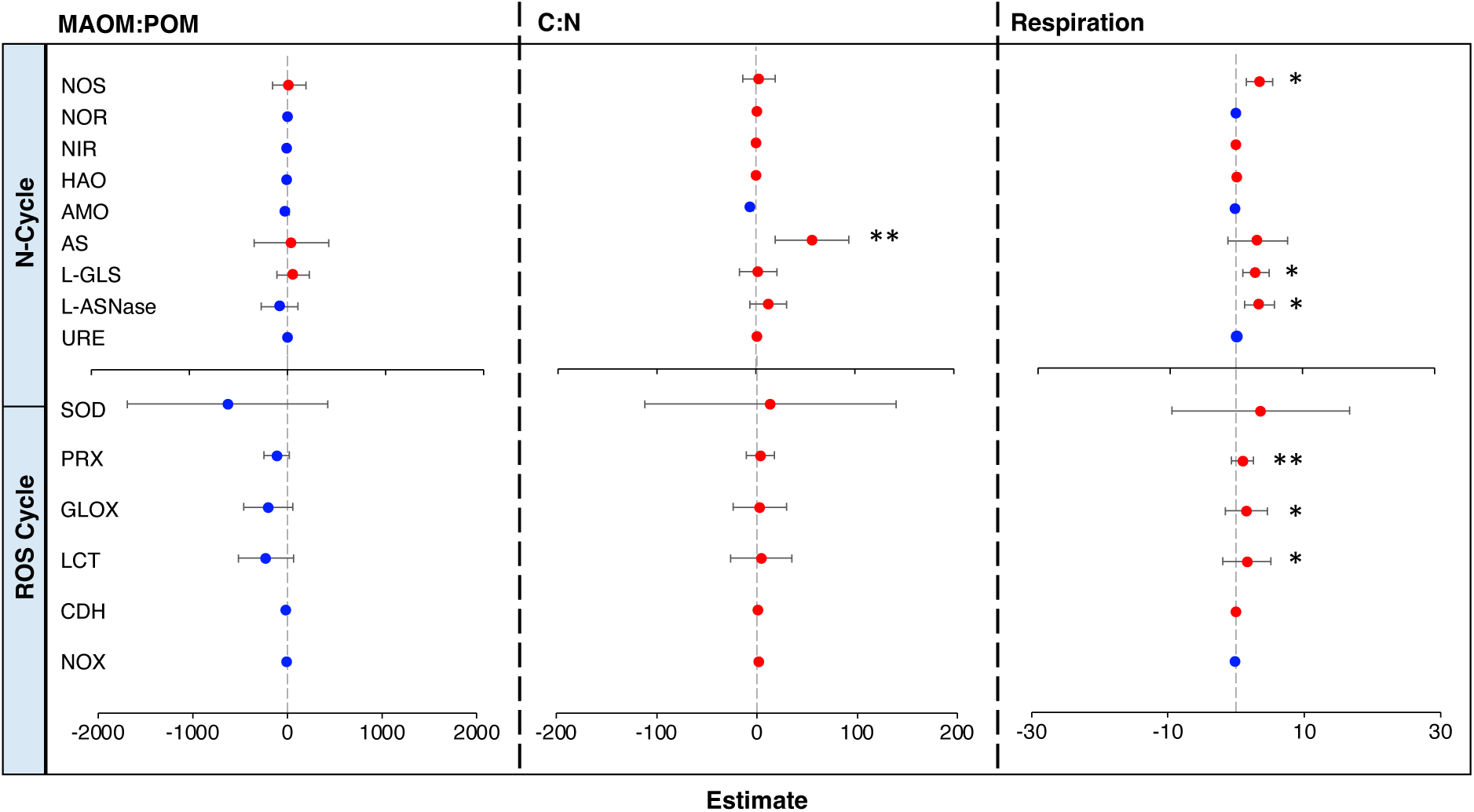
Mixed effects models. Effects of MAOM:POM, soil C:N, and microbial respiration N-cycle and ROS metatranscriptome data. Random effects were land-use type (“agriculture” “woodland”, and “built”) and human population size (“High Human Population [>400,000 Residents]” and “Low Human Population [<1000 Residents]”). Red indicates positive effects, blue indicates negative effects. Significance is indicated by * = *p* < 0.05, ** = *p* < 0.01. *N* = 18.

### 3.5 Microbial Respiration Rates and Relationship to N-cycle and ROS Metatranscriptomes

Respiration rate is described as mg-CO_2_ kg-soil^-1^ day^-1^. Respiration rates of samples were significantly positively associated with high human population sites (KW, *p* < 0.05) (**Fig. S3A**). Respiration rate was significantly positively associated with MAOM:POM ratio (SRCC, *p* < 0.05) (**Fig. S3B**). Mixed effects modelling shows positive associations between respiration rate and metatranscriptomic data for NOS, AS, L-GLS, and L-ASNase. There were also positive associations between respiration rate and the metatranscriptomes of ROS-cycling enzymes SOD, PRX, GLOX, and LCT. Potential net rates of ammonification were significantly higher in soils from locations with high human populations (KW, *p* < 0.01).

### 3.6 Partial Least Squares Path Models

Partial Least Squares Path Modelling (PLS-PM) showed positive effects of N-cycling metatranscriptomes, ROS-producing metatranscriptomes, soil processes, and soil properties on *F_NOy_* (effect size = 0.598, *p* < 0.05; effect size = 0.106, *p* > 0.05; effect size = 0.521, *p* < 0.05; effect size = 0.524, *p* < 0.05), respectively) (**Fig. 5**), of which N-cycling metatranscriptomes, soil processes, and soil properties were statistically significant effects (*p* < 0.05, for all). Human population and ROS-consuming metatranscriptomes had negative effects on *F_NOy_* (effect size = -0.201, *p* > 0.05; effect size = -0.242, *p* > 0.05, respectively).

**Figure 5.**
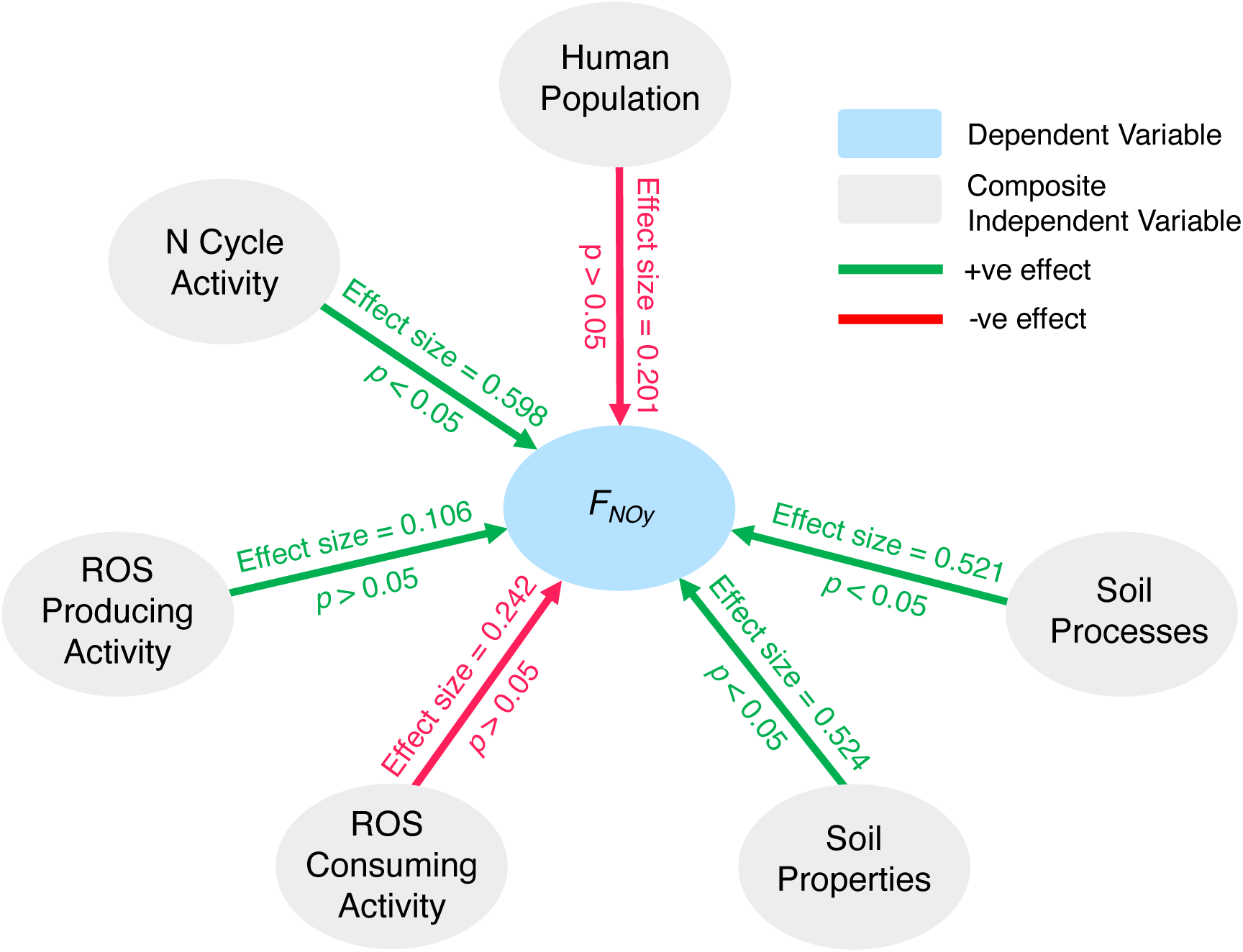
Partial Least Squares Path Modelling (PLS-PM) to ascertain effects of measured variables on fluxes of NO_y_ (*F_NOy_*). Fixed effects: human influence (human population size); composite N cycle activity (metatranscriptome of URE, L-ASNase, L-GLS, AS, AMO, HAO, NIR, NOR, and NOS); composite ROS producing activity ( metatranscriptome of NOX, CDH, and GLOX); composite ROS consuming activity (metatranscriptome of PRX, and SOD); composite soil properties (MAOM:POM, moisture content, available NO_3-_, available NH_4+_, total soil C:N); composite soil processes (net nitrification rate, net ammonification rate, total N mineralisation rate, respiration rate). *N* = 18.

## 4. Discussion

A landscape scale study of soil- sourced NO_y_ fluxes previously demonstrated large-scale heterogeneity in composition and processes of microbial communities involved with the N-cycle.^18^ The microbial communities of the soils examined in this work were distinct between land-uses and locations with different human populations, which we report as a function of soil properties and organic matter content, and link to differences in our measurements of potential NO_y_ fluxes. However, soil-sourced NO_y_ fluxes result from many abiotic and biotic mechanisms, of which many remain elusive.^18,41^ By coupling metagenomic and metatranscriptomic methods, we have developed possible links between ROS and N cycling that result in enhanced fluxes of NO_y_.

Links between nutrient cycles in soils, such as N and C, have been well studied. The cycling of ROS and N is inextricably linked in physiological systems where O_2•−_ reacts at near diffusion-controlled rate with NO to form peroxynitrite (ONOO^-^).^31,42^ ONOO^-^ can subsequently interact chemically with soil-prevalent compounds, including CO_2_, to form NO_2_ and NO_2_ˉ/HONO.^32^ Physiologically, this is a mechanism for lowering O_2•−_ concentration and NO bioactivity^43,44^, but has not been investigated in soil and other environmental systems. In this study we found that a common reaction involving NO and O_2•−_ may occur in soil systems, with ammonia-oxidising microbes producing NO as part of the N-cycle, and heterotrophic microbes producing O_2•−_ during the degradation of SOM. **Fig. 6** shows a proposed mechanism involving the interactions of ROS and N-cycling that may lead to enhanced NO_y_ emissions. We suggest that SOM composition and C:N determine the dominant taxa of heterotrophic microbes that decompose OM and make N available to other microbes, driving nitrification. There are several routes of NH_4+_ and NH_2_OH oxidation to NO_2-_ before it is reduced to NO by NIR. NO can be denitrified to N_2_ via N_2_O by NOR and NOS, but in this model there is excess N from anthropogenically enhanced N deposition that is likely lost to the atmosphere as described in the “leaky pipe” model of the N-cycle. ^45,46^ *F_NOy_* were also associated with increased activity of enzymes involved in organic matter decomposition and ROS production. When the ROS concentration is higher, ROS can react with NO produced during the N-cycle to form another NO_y_ species, NO_2_, via the intermediate ONOO^-^, potentially increasing NO_y_ fluxes. Our study presents evidence of rapid ONOO^-^ decomposition to NO_2_ in soils and indicates this NO_2_ is released to the atmosphere rather than reacting on heterogeneous surfaces in soil. Although the mechanism and kinetics of ONOOˉ decomposition has not been studied, it is likely dependent on soil conditions, including pH and temperature. We propose ONOOˉ decomposition proceeds via the pathways shown in reactions (R1) - (R2), two of which give rise to NO_y_ species.^47^ In addition to reaction (R1), it is possible that NO_2_ is also formed in the presence of CO_2_, which promotes the formation of an unstable product between ONOOˉ and CO_2_, which decomposes in part to NO_2_.^32^

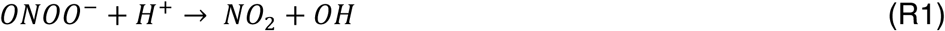

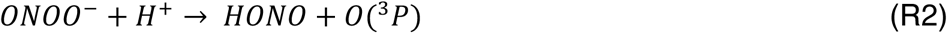

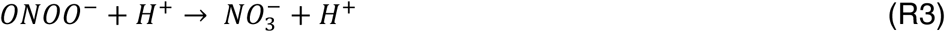

ONOOˉ is known to have a short half-life in physiological systems, which may translate to environmental systems. In the presence of reducing agents, such as Fe^2+^, the NO_2_ produced by ONOOˉ decomposition can also be reduced to NO_2_ˉ, which could then lead to emissions of NO_y_ species including HONO.^48,49^

**Figure 6.**
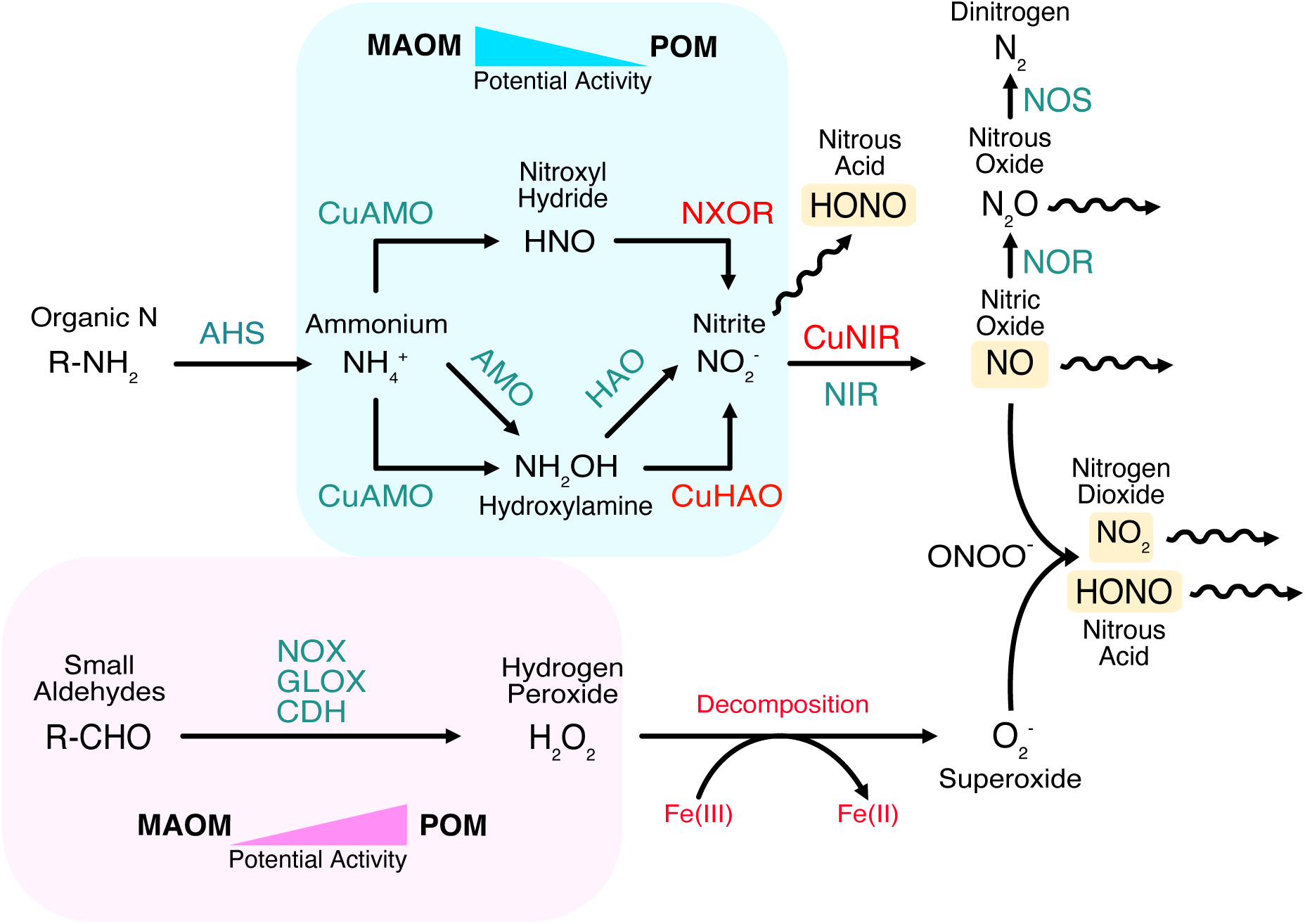
Proposed mechanism of soil sourced NOy production. Enzymes and processes in red are currently unresolved steps in the mechanism. Yellow boxes denote NOy species.

### 4.2 SOM Composition Determines Routes of Microbial Mineralisation

In this study, metatranscriptomes of ROS-producing and N-cycling enzymes were significantly positively correlated with *F_NOy_*, providing evidence for interactions between ROS and N species leading to NO_y_ production. Enzymes responsible for ROS scavenging such as SOD were associated with lower NO_y_ fluxes, further suggesting a link between ROS and N-cycling in the environment. As the microbial community can be significantly affected by SOM^50^, we suggest SOM composition determines routes of cycling of soil C and N. It has been reported that increased N deposition may contribute to an increase in the amount of organic matter stored as MAOM^51^, which we suggest is a contributing factor to the high MAOM:POM measured from more anthropogenically influenced soils in this study. Metatranscriptomes of amidohydralases AS, L-GLS, and L-ASNase were positively associated with respiration rate and MAOM:POM, suggesting this to be the primary N mineralisation route in MAOM dominant soils. After this mineralisation of organic N to NH_4+_, we suggest further N-cycling by autotrophic microbes is favoured and further aided by the steady nutrient supply and stable redox conditions in MAOM dominant soils, consequently leading to enhanced NO_y_ fluxes and outgassing. Metatranscriptomes of ROS-cycling enzymes were negatively associated with MAOM:POM. We suggest that in higher POM soils, heterotrophic microbial taxa that produce ROS as by-products of decomposition are more active. Increased concentrations of ROS can react with NO and prevent it from being emitted from soils.

POM has been described as a readily available source of C and N in soils due to its increased physical accessibility to microbial decomposers and plants compared to MAOM, and the rapid response time of this fraction to changes in soil nutrient levels or additions.^52^ The potential for faster decomposition of this fraction and resulting short lived nutrient pulses, particularly of CO_2_, may create conditions that favour ROS-producing heterotrophs^53,54^ which are more able to adapt to the dynamic environment. Whether autotrophic or heterotrophic pathways in soil dominate is dependent largely on redox conditions.^55^ As POM is decomposed, oxygen is consumed, and CO_2_ is produced via microbial respiration, which can create anaerobic microsites within the overall aerobic soil.^56^ Many heterotrophs respire aerobically or anaerobically depending on the O_2_ availability and these facultative heterotrophs may take advantage of the dynamic environment created by POM decomposition. Mineral- stabilised SOM is adsorbed to minerals and is less available than mineral-associated SOM.^57^ Whether OM adsorption to minerals reaches a point of saturation has been a topic of some debate, though it is likely that more robust methods of SOM fractionation would be required to define the saturation point conclusively.^58,59^ Although MAOM is often a longer-term sink for nutrients due to the protection from decomposition conferred by association with minerals, MAOM becomes available to microbes when it dissociates from minerals^60,61^ – likely much more so than POM. Therefore there is a proportion of MAOM with more available nutrients than may be first thought.^60^ This, and the higher C:N of this fraction as measured in this work, provides a steadier and more consistent release of labile N that may support the mineralising activity of chemoautotrophic bacteria. MAOM dominant soils contain smaller and denser particles^62^, which could lead to fewer and smaller soil pores with improved aggregation. This soil structure facilitates stable conditions in terms of oxygen and moisture availability, as well as maintaining a sufficient nutrient supply^63^, which would favour N-cycle processes (e.g., nitrification, etc.) in MAOM dominant soils. A previous study on soils from the same sites quantified heavy metal concentration, finding a lower concentration of Fe in high MAOM soils.^18^ This suggests less of this MAOM is adsorbed to Fe, and potentially other minerals in these soils, and therefore is more accessible for microbial mineralisation. Recent research has suggested that MAOM can be directly mobilised from mineral surfaces by some root exudates, such as oxalic acid. Other root exudates can also indirectly lead to MAOM mobilisation by stimulation of enzymatic activity.^64^ MAOM content tends to be a function of mineralogy, whereas POM content is due to vegetation cover, making location and land-use important factors in SOM composition.^65^ Plant cover differs between land-use types and the types and concentrations of root exudates will also differ. In woodland sample sites from this study, there was a diverse range of plant species growing which could have stimulated the release of N from the dominant MAOM fraction of these soils. Plant composition is likely to influence SOM dynamics, nutrient and mineral content of soil, and microbial community composition. Direct plant influences on soil, through roots and litter properties, and the consequences for this proposed mechanism will need to be addressed in future work.

### 4.3 Pathways to NO_y_ by Ammonia Oxidisers

As discussed above, in soils from high human population sites, the MAOM fraction of SOM dominates. N is available to the microbial community in these soils because it has dissociated from the minerals, either through saturation of mineral adsorption or mobilisation by microbial activity or root exudates. Processes of the N-cycle, including the formation of NO_y_, are enhanced by the increased availability of N in these soils. After organic N has been mineralised to form NH_4+_ there are several potential pathways where NH_4+_ is converted to NO via ammonia oxidation. Although ammonia oxidising bacteria (AOB) and archaea (AOA) can coexist, differences in their ecological niches may lead to competition.^66^ Higher concentrations of NH_4+_ are expected in these soils due to enhanced mineralisation as a consequence of excess N deposition and AOB are more competitive against AOA under these high N conditions.^67^ Varying relative abundance of AOA and AOB communities is closely correlated with changing soil pH and AOA may be more resilient to fluctuating O_2_ availability.^68,69^ The highly dynamic soil environment means the dominant ammonia oxidising taxa often changes depending on conditions.

In this work, the abundance of microbes with the functional potential for ammonia oxidation was positively correlated with *F_NO_*, suggesting nitrifier denitrification as a possible mechanism for NO production. However, AOB taxa are able to further denitrify NO to N_2_O and N_2_ and so we would not expect to see high emissions of *F_NO_* from soils, particularly in high MAOM soils where denitrification may be enhanced by the likely reducing conditions. Production of NO_y_ is primarily a result of conventional nitrification and denitrification. However, under certain conditions, nitrifier denitrification can also contribute significantly.^70,71^ Ammonia oxidisers have been found to be responsible for up to 56% of N_2_O emissions from soils^71^, however relative contribution of nitrifier denitrification to NO emissions has yet to be conclusively quantified. Nitrifier denitrification is more common under high N conditions as there is more availability of substrates including NH_4+_ and NO_3-_ that stimulate the activity of ammonia oxidising taxa. While highly aerobic soils favour nitrification, nitrifier denitrification can be enhanced when less O_2_ is available such as when moisture content is high or soil is made up of smaller and denser particles – as is seen in high MAOM:POM soils.^62^ Respiration rate is often higher in soils with high N inputs due to increased microbial activity as was the case in this work, which would further contribute to the depletion of O_2_ in these soils. Increased metatranscriptomic reads of amoA encoded by *Nitrososphaerales*, an AOA taxon, was significantly associated with soils from which high MAOM:POM was measured. However, the capability of AOA to carry out nitrifier denitrification has been hypothesised, but not experimentally proven. As such, it is difficult to conclude that increased *F_NO_* in these soils is due to nitrifier denitrification, however conditions are suitable to favour this process and could be possible.

During ammonia oxidation, NH_2_OH is produced by AMO. In AOB, NH_2_OH is then oxidised to NO_2-_ by HAO and NO_2-_ can then be converted to NO by NIR as part of the nitrifier denitrification pathway. AOA lack the HAO enzyme, however they do encode Cu-containing periplasmic oxidase proteins, of which one could act as an hydroxylamine oxidiser, CuHAO.^72,73^ It is possible that AOA do not produce NH_2_OH during ammonia oxidation, and instead encode a different AMO that produces the highly reactive and unstable intermediate nitroxyl hydride (HNO). HNO can be a direct source of N_2_O by dimerization to hyponitrous acid (H_2_N_2_O_2_) and subsequent dehydration.^74^ NO_2-_ could be produced from HNO via one of the Cu-containing periplasmic oxidase proteins acting as a nitroxyl oxidoreductase (NXOR). From the CuHAO or NXOR routes discussed above, NO can be formed following nitrite production. It has been reported that *Nitrosopumilus maritimus*, a marine AOA, has genes encoding a Cu-containing NirK.^72^ Whether AOA produce NO via the CuHAO or the NXOR pathway would likely be determined by which of their encoded AMO enzymes has a higher affinity for NH_4+_. The multiple pathways of N utilisation in AOA may be advantageous, as they are able to utilise available N more rapidly and diversely than AOB. In contrast to AOB, which rely on Fe-containing enzymes, AOA seem to utilise Cu-containing enzymes. We hypothesise therefore that in soils with higher Cu concentrations, AOA may be more competitive than AOB, and contribute more to soil-sourced *F_NOy_*. In previous work^18^, Fe concentrations were found to be negatively correlated with soils from high human population locations, which had high MAOM:POM here, suggesting AOA may outcompete Fe-reliant AOB and lead to increased *F_NOy_*.

### 4.4 Nitric Oxide Emitted from the “Leaky Pipe” Model

Metatranscriptome counts of genes encoding NIR, NOR and NOS were positively correlated with NO_y_, suggesting a higher reductive potential in soils with higher MAOM. However this is a process that converts NO to other N species, N_2_O and N_2_ and would be expected to limit NO loss to the atmosphere. The “leaky pipe” model imagines the N-cycle as N flowing through pathways with inefficiencies through which N species, particularly NO and N_2_O, can be lost.^45,46^ In this model, the loss of NO from the N-cycle is dependent on the rates and efficiency of nitrification and denitrification, which are in turn affected by how much N is available in the soil, the soil moisture content, and soil redox conditions. Excess N inputs exacerbate the leaks in the pipe by driving N-cycle processes of NO production and in the case of these aerobic soils, much of the NO produced is lost from soils before it can be denitrified further by anaerobic denitrifiers.^45^ Gases such as NO that have been lost from the N-cycle can be taken up by plants, by microbes in soil pores, or oxidised to NO_2_ (60- 95%)^75–77^, but there are also several other possible outcomes^46^. MAOM dominant soils contain smaller and denser particles^62^, which could lead to fewer and smaller soil pores, and therefore less opportunity for NO to be taken up rather than emitted, though this is yet to be experimentally proven. In the context of this study, we suggest that anthropogenic N deposition to soils provides sufficient N that the NO produced by nitrification can be denitrified, or if it is lost from the N-cycle can react with ROS to form NO_2_ or be emitted to the atmosphere. Much research has been undertaken to quantify losses of NO and N_2_O from agricultural soils after fertiliser addition, and devise actions to improve fertiliser efficiency, enhance crop yields, and minimise environmental impacts.^78–80^ However, consequences of the “leaky pipe” model for NO_y_ emissions have yet to be fully quantified. Interactions between the ROS and N-cycles examined in this work may shed light on further routes to soil-sourced gas emissions that must be considered.

## 5. Conclusions

As a result of this work, we suggest that ROS and N-cycling in soils is inextricably linked with consequences for NO_y_ fluxes in these systems. While our previous work identified spatiotemporal patterns in soil NOy emissions on a landscape scale, this study reveals a novel mechanistic pathway linking ROS and N cycling through ONOO^-^ chemistry, providing new perspective on both biotic and abiotic controls of soil trace gas production. PLS-PM showed significant positive effects of metatranscriptomes of both ROS and N-cycling genes on *F_NOy_*. Reactions between ROS and N species that are prevalent in physiological systems seem to translate to soil systems and lead to the formation of NO_y_ species. Soil properties, particularly the composition of SOM, affects how significantly these ROS mechanisms contribute to NO_y_ fluxes relative to more widely investigated N-cycle processes. Future research should explore how this pathway responds to global change drivers, particularly as climate change alters soil moisture and temperature patterns, and organic matter dynamics. The mechanism proposed in this study opens new avenues for understanding soil-atmosphere interactions and their role in global biogeochemical cycles. By coupling metagenomic and metatranscriptomic data we have demonstrated that relying on metagenomic data alone, which only captures the potential for microbial processes rather than activity, could mean vital mechanistic data for microbial dynamics is missed. The interaction between ROS and N cycling revealed here has important implications for biogeochemical modelling. Current terrestrial ecosystem models typically treat N cycling and oxidative processes separately, potentially missing important feedbacks that influence trace gas emissions. Incorporating these interactions could improve predictions of soil NO_y_ fluxes across scales, from ecosystem to global levels. This is particularly relevant for understanding how land-use change and urbanisation might alter soil-atmosphere gas exchange. Long-term studies across diverse ecosystems would help quantify the relative contribution of this pathway to global NO_y_ budgets and its sensitivity to environmental change. Additionally, investigation of how this mechanism varies seasonally and across climatic gradients would improve our ability to predict and manage soil NO_y_ emissions in a changing world.

## Data Availability

Data are available at Purchase, Megan L (2024), “Dynamics of Reactive Oxygen Species and Nitrogen Cycling in Soils as a Mechanism for Volatile Reactive Nitrogen Oxide Production”, Mendeley Data, V1, doi: 10.17632/kj6x2b4zrg.1 Raw sequence data have been deposited in the NCBI Sequence Read Archive under accession number PRJNA1185162. Code is available at https://github.com/MeganPurchase/Dynamics_ROS_NOy_Purchase24.git

## Acknowledgements

Financial support for this study came from the UKRI Natural Environment Research Council (NERC) as part of the Central England NERC Training Alliance (CENTA2) grant NE/S007350/1, and the UKRI Biotechnology and Biological Sciences Research Council (BBSRC) as part of grant BB/X002187/1. Sequencing was funded as a pilot project by the NERC environmental omics facility (NEOF1489). The authors thank members of the Mushinski lab group for feedback on this manuscript.

