## Supporting Information for "Dynamics of Reactive Oxygen Species and Nitrogen Cycling in Soils as a Mechanism for Volatile Reactive Nitrogen Oxide Production"

**Table of Contents**

**Methods**

Method S1. NOy Flux Calculations……………………………………………2

Method S2. Soil Texture Calculations……………………………..…………..2

Method S3. Soil Respiration Calculations……...………………………………2

Method S4. N Mineralisation Assay Calculations………..…..………………...3

**Tables**

Table S1. Measured potential NOy, NO, and $NO_2$ + NOz fluxes………………4

Table S2. Soil Texture Classifications…………………………………………..5

**Figures**

Figure S1. Map of Sampling Sites………………………………………………6

Figure S2. Analysis of Soil Properties…………...……………………………..7

Figure S3. Analysis of Soil Processes…………………………..…….……….....8

**Methods**

**S1. NOy Flux Calculations**

Fluxes ($F_{soil}$) of NOy, NO, and $NO_2$+NOz (ng-N $m^{-3}$ $hr^{-1}$) were calculated as below:

$$F_{soil} = (Conc._{soil} - Conc._{blank})x\ flow\ x\ (\frac{Chamber\ volume}{Soil\ surface\ area})$$

**S2. Soil Texture Calculations**

Relative proportions of sand, silt, and clay (% $L^{-1}$) were calculated from hydrometer and temperature readings.

1. For each degree below 20°C, 0.36 g L-1 was subtracted from the hydrometer reading.
2. To calculate the silt + clay fraction, the corrected blank hydrometer reading was subtracted from the corrected sample readings at 40 seconds. The resulting value was divided by the sample weight then multiplied by 100 to give silt + clay % $L^{-1}$.
3. To calculate the clay fraction, the corrected blank hydrometer reading was subtracted from the corrected sample readings at 2 hrs. The resulting value was divided by the sample weight then multiplied by 100 to give clay % $L^{-1}$.
4. The silt fraction was calculated by subtracting clay % $L^{-1}$ from silt + clay % $L^{-1}$.
5. The sand fraction was calculated by subtracting silt + clay % $L^{-1}$ from 100.
6. Textural classes were assigned using **Figure S2**.

**S3. Soil Respiration Calculations**

Dry soil mass ($m_{Bt}$; in kg) was calculated:

$$m_{Bt} = m_{Bf} * \frac{TS}{100}$$

$m_{Bf}$ = wet soil mass (kg)

$TS$ = dry soil content (%)

Free gas volume ($V_{fr}$; in L) was calculated:

$$V_{fr} = V_{ges} - V_{AG} - V_{AM} - V_{Bf}$$

$V_{ges}$ = total volume of headspace enclosed in the measuring vessel – without soil, absorption vessel or absorbing agent (L)

$V_{AG}$ = volume of the absorption vessel (L)

$V_{AM}$ = volume of the absorbing agent (L)

$V_{Bf}$ = wet soil volume (L)

Soil respiration (mg-$O_2$-consumed/kg-soil) was calculated:

$$Soil\ respiration = \frac{Mr(O_2)}{R * T} * \frac{free\ gas\ volume\ (L)}{dry\ soil\ mass} * \Delta p$$

$Mr(O_2)$ = molar mass of $O_2$ (32000 mg/mol)

$R$ = general gas constant (83.14 L mbar $mol^{-1}$ $K^{-1}$)

$T$ = measuring temperature (K)

$\Delta p$ = reduction in pressure of the measuring preparation (mbar)

**S4. N Mineralisation Assay Calculations**

Concentrations were converted from a mass nutrient per unit volume to a mass nutrient per mass dry soil weight using the below calculations:

$$\frac{[NO_3^-]mg}{L} * \frac{extraction\ vol.(L)}{soil\ wet\ weight\ (g) * (dry\ weight\colon wet\ weight)} * \frac{1000\mu g}{1\ mg} = [NO_3^-]\frac{\mu g}{g\ dw}$$

$$\frac{[NH_4^+]mg}{L} * \frac{extraction\ vol.(L)}{soil\ wet\ weight\ (g) * (dry\ weight\colon wet\ weight)} * \frac{1000\mu g}{1\ mg} = [NH_4^+]\frac{\mu g}{g\ dw}$$

Net nitrification and net ammonification rates were calculated:

$$\frac{([NO_3^- \frac{\mu g}{g\ dw}]_{final} - [NO_3^- \frac{\mu g}{g\ dw}]_{initial})}{incubation\ time\ (days)} = Nitrification\ rate(\mu g\ NO_3^- \cdot g^{-1} \cdot day^{-1})$$

$$\frac{([NH_4^+ \frac{\mu g}{g\ dw}]_{final} - [NH_4^+ \frac{\mu g}{g\ dw}]_{initial})}{incubation\ time\ (days)} = Ammonification\ rate(\mu g\ NH_4^+\ \cdot g^{-1}\ \cdot day^{-1})$$

N Mineralisation rates were calculated:

$$\frac{(\Sigma\ inorg.N_{final} - \ \Sigma\ inorg.N_{initial})}{incubation\ time\ (days)} = N\ min.rate\ (\mu g\ inorganic\ N\ \cdot g^{-1}\ \cdot\ day^{-1})$$

**Tables**

**Table S1. Fluxes of NOy, NO, and $NO_2$+NOz (ng-N $m^{-3}$ $hr^{-1}$)**

| | | Flux (ng-N $m^{-3}$ $hr^{-1}$) | | |
|---|---|---|---|---|
| Location | Land-use Type | NOy | NO | $NO_2$ +NOz |
| **Coventry** | Agriculture | 417.82 | 453.08 | -34.78 |
| **Coventry** | Agriculture | -287.56 | 41.13 | -328.53 |
| **Coventry** | Agriculture | -11.21 | 152.81 | -164.03 |
| **Coventry** | Woodland | 632.21 | 715.67 | -83.47 |
| **Coventry** | Woodland | 4888.61 | 4892.23 | -3.86 |
| **Coventry** | Built | 98.13 | 128.40 | 3.55 |
| **Coventry** | Built | 428.85 | 406.32 | 22.51 |
| **Coventry** | Built | 378.89 | 418.10 | -39.21 |
| **Wellesbourne** | Agriculture | 1400.94 | 1103.09 | 297.79 |
| **Wellesbourne** | Agriculture | -124.89 | 76.26 | -211.12 |
| **Wellesbourne** | Agriculture | 182.63 | 259.96 | -77.32 |
| **Wellesbourne** | Woodland | -532.45 | 103.35 | -635.44 |
| **Wellesbourne** | Woodland | -87.22 | 32.853 | -120.09 |
| **Wellesbourne** | Woodland | -596.28 | 1851.68 | -1851.68 |
| **Wellesbourne** | Built | 100.55 | 75.764 | 24.91 |

| | | | | |
|---|---|---|---|---|
| **Wellesbourne** | Built | -79.28 | 121.45 | -200.75 |
| **Wellesbourne** | Built | 1320.36 | 1237.98 | 82.33 |

**Table S2. Soil Texture Classifications**

| Location | Land-use Type | Soil Texture | Clay % | Silt % | Sand % |
|---|---|---|---|---|---|
| **Coventry** | Agriculture | Sandy loam | 12.503 | 20.005 | 67.492 |
| **Coventry** | Agriculture | Sandy loam | 14.996 | 14.996 | 70.007 |
| **Coventry** | Agriculture | Loamy sand | 14.981 | 17.478 | 67.541 |
| **Coventry** | Woodland | Sandy loam | 2.503 | 15.019 | 82.478 |
| **Coventry** | Woodland | Sandy loam | 4.996 | 19.985 | 75.019 |
| **Coventry** | Woodland | Sandy loam | 19.990 | 2.499 | 77.511 |
| **Coventry** | Built | Sandy loam | 2.503 | 12.513 | 84.985 |
| **Coventry** | Built | Sandy loam | 5.001 | 22.506 | 72.493 |
| **Coventry** | Built | Sandy loam | 5.003 | 32.516 | 62.481 |
| **Wellesbourne** | Agriculture | Sandy loam | 12.494 | 24.988 | 62.519 |
| **Wellesbourne** | Agriculture | Sandy loam | 15.789 | 28.947 | 55.263 |
| **Wellesbourne** | Agriculture | Sandy loam | 7.509 | 10.013 | 82.478 |
| **Wellesbourne** | Woodland | Loamy sand | 15.004 | 17.504 | 67.492 |
| **Wellesbourne** | Woodland | Loamy sand | 10.000 | 15.000 | 75.000 |
| **Wellesbourne** | Woodland | Sandy clay loam/Sandy loam | 17.491 | 19.990 | 62.519 |
| **Wellesbourne** | Built | Loamy sand | 15.008 | 20.010 | 64.982 |
| **Wellesbourne** | Built | Sandy loam | 17.504 | 15.004 | 67.492 |
| **Wellesbourne** | Built | Sandy loam | 7.498 | 32.492 | 60.009 |

### Figures

#### Figure S1. Map of Sampling Sites

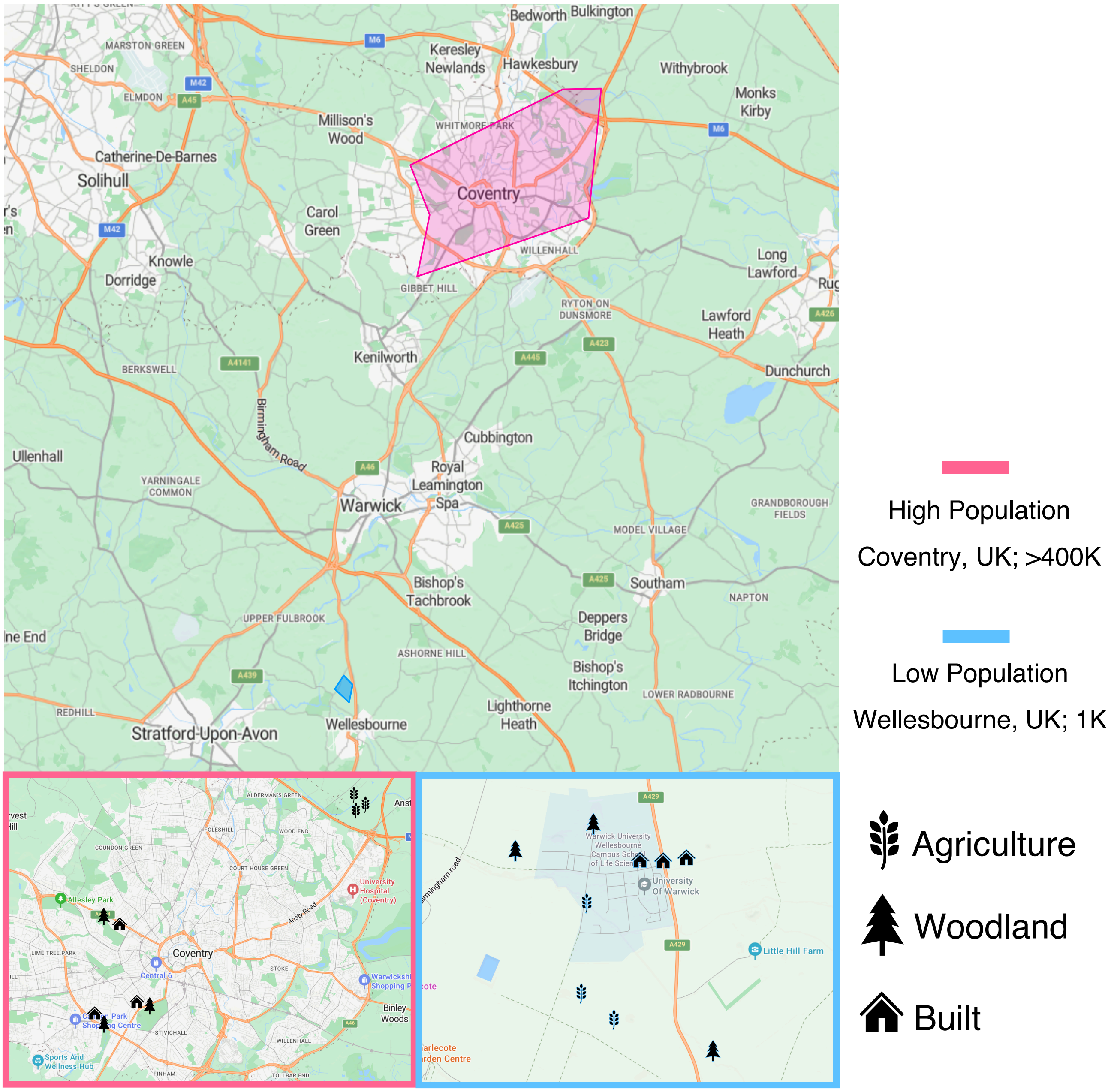


**Figure S1.** Map showing sampling sites in a high population area (Coventry, UK; >400K) and a low population area (Wellesbourne, UK; <1K). Markers indicate land-use types sampled ( = agriculture, = woodland, and = built).

**Figure S2. Analysis of Soil Properties**

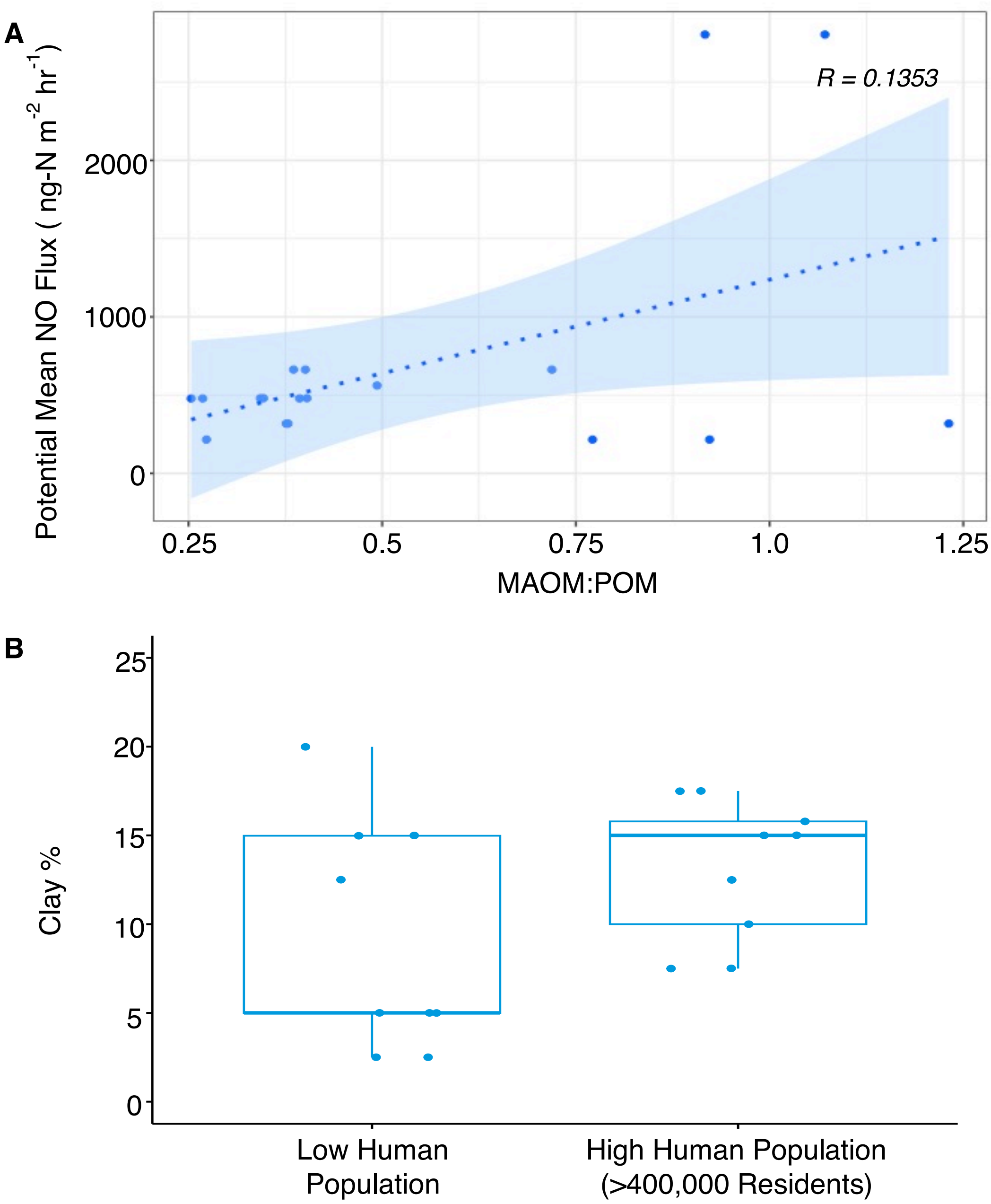


**Figure S2. (A)** SOM composition (MAOM:POM) plotted against potential mean fluxes of NO. Fluxes were measured with a chemiluminescence technique using a Teledyne T200U instrument. *N* = 18. **(B)** Clay % of soil, quantified using size fractionation, plotted against human population ("High Human Population [>400,000 Residents]", "Low Human Population [<1000 Residents]"). *N* = 18.

**Figure S3. Analysis of Soil Processes**

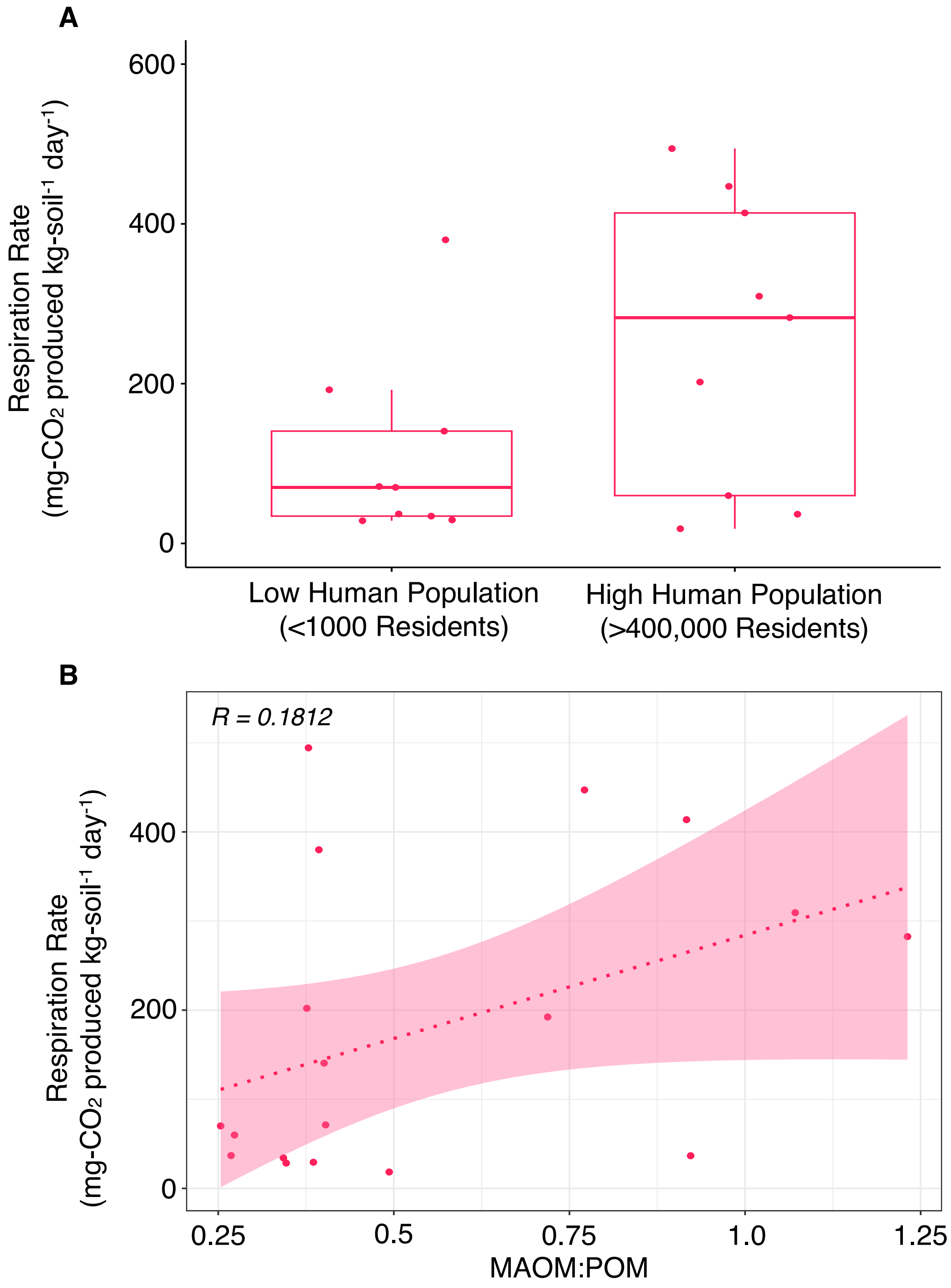


**Figure S3. (A)** Respiration Rate (mg-$CO_2$ produced kg-soil$^{-1}$ day$^{-1}$) plotted against human population ("High Human Population [>400,000 Residents]", "Low Human Population [<1000 Residents]"). *N* = 18. **(B)** Respiration Rate (mg-$CO_2$ produced kg-soil$^{-1}$ day$^{-1}$), plotted against SOM composition (MAOM:POM). Respiration rates were measured using the OxiTop® respirometric BOD measuring system. *N* = 18.
